# Identification of Six Thiolases and their Effects on Fatty Acid and Ergosterol Biosynthesis in *Aspergillus oryzae*

**DOI:** 10.1101/2021.12.02.471048

**Authors:** Hui Huang, Yali Niu, Qi Jin, Kunhai Qin, Li Wang, Bin Zeng, Zhihong Hu

## Abstract

Thiolase plays important roles in lipid metabolism. It can be divided into degradative thiolases (Thioase I) and biosynthetic thiolases (thiolases II), which are involved in fatty acid β-oxidation and acetoacetyl-CoA biosynthesis, respectively. The *Saccharomyces cerevisiae* (*S. cerevisiae*) genome harbors only one gene each for thioase I and thiolase II, namely, *Pot1* and *Erg10*, respectively. In this study, six thiolases (named AoErg10A−AoErg10F) were identified in *Aspergillus oryzae* (*A. oryzae*) genome using bioinformatics analysis. Quantitative reverse transcription–PCR (qRT-PCR) indicated that the expression of these six thiolases varied at different growth stages and under different forms of abiotic stress. Subcellular localization analysis showed that AoErg10A was located in the cytoplasm, AoErg10B and AoErg10C in the mitochondria, and AoErg10D-AoErg10F in the peroxisome. Yeast heterologous complementation assays revealed that AoErg10A, AoErg10D, AoErg10E, AoErg10F and cytoplasmic AoErg10B (AoErg10B^ΔMTS^) recovered the phenotypes of *S*. *cerevisiae erg10* weak and lethal mutants, and that only AoErg10D-F recovered the phenotype of the *pot1* mutant that cannot use oleic acid as the carbon source. Overexpression of AoErg10s either affected the growth speed or sporulation of the transgenic strains. In addition, the fatty acid and ergosterol content changed in all the AoErg10-overexpressing strains. This study revealed the function of six thiolases in *A. oryzae* and their effect on growth, and fatty acid and ergosterol biosynthesis, which may lay the foundation for genetic engineering for lipid metabolism in *A. oryzae* or other fungi.

**Importance:** Thiolase including thioase I and thiolase II, plays important roles in lipid metabolism. *A. oryzae,* one of the most industrially important filamentous fungi, has been widely used for manufacturing oriental fermented food such as sauce, miso, and sake for a long time. Besides, *A. oryzae* has a high capability in production of high lipid content and has been used for lipid production. Thus, it is very important to investiagte the function of thiolases in *A. oryzae*. In this study, six thiolase (named AoErg10A-AoErg10F) were identified by bioinformatics analysis. Unlike other reported thiolases in fungi, three of the six thiolases showed dual function of thioase I and thiolase II in *S. cerevisiae*, indicating the lipid metabolism is more complex in *A. oryzae*. The reveal of founction of these thiolases in *A. oryzae* can lay the foundation for genetic engineering for lipid metabolism in *A. oryzae* or other fungi.

## Introduction

Thiolases, also known as acetyl-coenzyme A acetyltransferases, are enzymes found abundantly in both prokaryotes and eukaryotes (1). Thiolases play important roles in lipid metabolism. Based on the direction of the reaction catalyzed, thiolases can be divided into two categories: degradative and biosynthetic thiolases (2). The former group, also known as thioase I, includes β- ketoacyl-CoA thiolases (KTs; EC 2.3.1.16), which catalyzes the removal of an acetyl group from a β-ketoacyl-CoA, yielding acetyl-CoA and shorter acyl CoA molecule. This catabolic reaction mainly occurs in the last step of every catalytic cycle of fatty acid β-oxidation. In contrast, the biosynthetic thiolases, also known as thiolase II, catalyze the condensation of two molecules of acetyl-CoA to form acetoacetyl-CoA. This reaction is the first enzymatic step in many anabolic processes such as the biosynthesis of sterols and ketone bodies in prokaryotes or the synthesis of poly (3-hydroxybutyric acid), a major energy and carbon storage molecule in many bacteria.

The substrate specificities of the two types of thiolases differ: thiolase I is specific for substrates 4 to 22 carbon atoms in length, while thiolase II is specific for C4 chains. Mostly, the two type thiolases can catalyze both biosynthetic and degradative reactions. Thiolase I can only effectively catalyze the degradative reaction, which is thermodynamically favorable; thiolase II can also act in the last thiolytic cleavage of the fatty acid β-oxidation spiral (3). Bioinformatics studies have shown that the two classes of thiolases originated from a common ancestor, as they share significant sequence similarity and essentially use the same active site residues to perform relevant reactions (1, 3). The two types of thiolases are mainly localized in three different compartments: peroxisome, mitochondria, and cytosol. Thiolase I is localized in the mitochondria and peroxisome, where it metabolizes fatty acids for energy production in eukaryotes. In contrast, in addition to biosynthesis of sterols or ketone bodies, thiolase II can also be found in animal mitochondrion (involved in the metabolism of ketone bodies and isoleucine) and plant or fungi peroxisomes (completes the degradation of fatty acids) (4–6). Some eukaryotic cells contain an additional peroxisomal thiolase I, known as SCP-X, which is specific for β-oxidation long- and branched-chain fatty acids. In some cases, this results as a fusion between a thiolase and a sterol carrier protein (7). Both types of thiolases function in the form of tight dimers or tetramers (dimers of tight dimers) (8).

Six different classes of thiolases (named CT, T1, T2, SCP2, AB and TFE) have been identified in humans; each of these thiolases differ from others with respect to sequence, oligomeric state, substrate specificity, and subcellular localization. Among these six thiolases, only CT-thiolase belongs to thiolase II and is located in the cytosol. SCP2 and AB are both peroxisomal degradative thiolases, and T1, T2, and TFE are mitochondrial degradative enzymes (9, 10). *Arabidopsis* contains nine thiolases, including four thiolase I (KAT1, KAT2, KAT5.1, and KAT5.2) and five thiolase II (ACAT1.1, ACAT1.2, ACAT1.3, ACAT2.1, and ACAT2.2). KAT1 and KAT2 are located in the peroxisome. KAT5.1 and KAT5.2 are encoded by different alternative spliced transcripts of the same gene. As the N-terminal sequences vary, KAT5.1 localizes in the cytosol, while KAT5.2 localizes to the peroxisome. Similarly, ACAT1.1, ACAT1.2, and ACAT1.3 are also encoded by the different alternative spliced transcripts of the same gene. ACAT1.1 and ACAT1.2 are cytosolic protein, while ACAT1.3 is a peroxisomal protein. ACAT2.1 and ACAT2.2 are also encoded by the same gene and localize in the cytosol (4). *S. cerevisiae* has only two thiolases. Thiolase I, also known as Pot1, localizes in the peroxisome and is involved in β-oxidation of fatty acids, and the pot mutant is reported to be lethal in medium with oleic acid as the sole carbon source (11, 12). Thiolase II, also known as Erg10, localizes in the cytosol and is involved in the first step of ergosterol biosynthesis where it condenses two acetyl CoA molecules to acetoacetyl coenzyme A (13). Similar to *S. cerevisiae*, *Xanthophyllomyces dendrorhous* harbors two thiolases (Erg10 and Pot1); *Erg10* is essential for ergosterol biosynthesis, while *Pot1* is not essential for cell growth, although it indirectly affects pigment production (14). Two thiolase II genes were identified in *Magnaporthe oryzae* (*MoAcat1* and *MoAcat2*); MoAcat1 and MoAcat2 showed different subcellular localization and promoted efficient development of cell morphology and effective colonization of host tissue, respectively (15). More recently, two thiolase II proteins, named AfErg10A and AfErg10B, were identified in *A. fumigatus*; AfErg10A is localized in the mitochondria and is essential for *A. fumigatus* survival and morphological development (16), while AfERG10B is essential for the viability of *A. fumigatus* (17).

However, studies on the function of thiolases in *A. oryzae,* one of the most industrially important filamentous fungi, are scarce. *A. oryzae* is a Food and Drug Administration- and World Health Organization-identified safe production filamentous fungus that has been widely used for manufacturing oriental fermented food such as sauce, miso, and sake for a long time. It is also commercially used as an excellent host for recombinant enzymes and secondary metabolite production due to its high secretion rates (18, 19). Unlike that reported for other fungi, our previous bioinformatics studies have shown six thiolase-encoding genes in the *A. oryzae* genome (20). Nevertheless, the function of these six thiolases is still unclear. In this study, six thiolases (named AoErg10A-AoErg10F) were identified using bioinformatics analysis, their expression pattern, subcellular localization, biochemistry founction and effects on fatty acid and ergosterol biosynthesis were detected. Therefore, this study revealed the function of six thiolases in *A. oryzae*, which will lay the foundation for genetic engineering of lipid metabolism in *A. oryzae* or other fungus.

## Materials and methods

### Phylogenetic Analysis

The unrooted tree was created in MEGA 6.0 using the neighbor-joining method with the following amino acid sequences. Cytoplasmic thiolase/acetyl-CoA C-acetyltransferase: *S. pombe* [Q9UQW6.1], *S. cerevisiae* [P41338.3], *B. graminis* [EPQ61678.1], *N. tabacum* [AAU95618.1], *A. thaliana* [Q9FIK7.1], *Z. mays* [NP_001266315.1]. Mitochondrial thiolase/acetyl-CoA C-acetyltransferase: G. gallus [NP_001264708.1], *R. norvegicus* [NP_058771.2], *H. sapiens* [BAA01387.1], *B.taurus* [NP_001039540.1], *C. lupus familiaris* [XP_546539.2]. Mitochondrial thiolase/3-ketoacyl-CoA thiolase: *B. taurus* [NP_001030419.1], *M. musculus* [NP_803421.1], *R. norvegicus* [NP_569117.1]. Peroxisomal thiolase/3-ketoacyl-CoA thiolase: *S. cerevisiae* [CAA37472.1], *Y. lipolytica* [Q05493.1], *A. thaliana* ([AEE27736.1], [AEC0879.1] and [AED95738.1]), *G. gallus* [NP_001184217.1], *M. musculus* [NP_570934.1], *H. sapiens* [NP_001598.1], *B. taurus* [NP_001029491.1], *E. caballus* [XP_001488609.1], *C. lupus familiaris* [XP_534222.2]. The accession number of *A. oryzae* thiolases, AoErg10A to AoErg10F, are as follows: EIT73661, EIT73496, EIT79671, EIT78121, EIT78942 and EIT80840. The accession numbers are provided in parentheses.

### Strains and Growth Conditions

*A. oryzae* 3.042 (CICC 40092), obtained from the China Center of Industry Culture Collection (Beijing, China), was used as the wild type strain. The uridine/uracil auxotrophic (△*pyrG*) *A. oryzae* 3.042 strain was constructed in our laboratory (19). *A. oryzae* was cultured on Czapek-Dox Agar medium (2% sucrose, 0.2% NaNO_3_, 0.1% KH_2_PO_4_, 0.05% MgSO_4_, 0.05% KCl, 0.05% NaCl, 0.002% FeSO_4_, 1.6% agar, pH 5.5) at 30°C. Spores were harvested after 3 days of culture and then suspended in sterile water containing 0.05% Tween-80. The concentration of spores was determined using a hemocytometer. *Escherichia coli* DH5α (21) was used for plasmid construction and *Agrobacterium tumefaciens* AGL1 (22) was used for *A. tumefaciens*-mediated transformation of *A. oryzae*. Both *E. coli* and *A. tumefaciens* were cultured in Luria Bertani (LB) medium supplemented with suitable antibiotics at 37°C and 28°C, respectively.

### Gene Expression Analysis

The mycelia at different growth stages or under different stress conditions were harvested and immediately frozen in liquid nitrogen and pulverized. Total RNA was isolated using a fungal RNA kit (Omega Bio-tek, Norcross, GA, USA) according to the manufacturer’s instructions after RNase-free DNase I treatment (Omega). The quality and concentration were determined using a NanoDrop ND-2000 spectrophotometer (Thermo Scientific, Wilmington, DE, USA). The cDNAs were synthesized from 1 μg total RNA using the Prime Script™ RT reagent kit (Perfect Real Time; Takara). All qRT–PCR (quantitative reverse transcription–PCR) operations were performed using a CFX96 real-time PCR detection system (Bio-Rad, CA, USA) using SYBR Premix Ex Taq (Takara, Japan). The housekeeping gene encoding histone H4 was used as the normalization control (23) and the relative expression was calculated using the formula 2^−ΔΔCt^. The sequences of the primers used for qRT–PCR are shown in Table S1.

### Functional Complementation in Yeast

The *erg10* (Y40985 and Y22800) and *pot1* (Y02319) mutants were purchased from EUROSCARF (http://www.euroscarf.de/index.php) and the corresponding yeast strain, BY4741/BY4743, were used as the wild-type control. Plasmid pYES 2.0 was used as the vector for yeast complementation. The full-length coding sequence (CDS) of *AoErg10s* and *Pot1* were fused into pYES 2.0 using a one-step cloning kit (Vazyme Biotech Co., Ltd, China). Then, the constructed vectors were transformed into the corresponding yeast mutant using the yeast transformation kit II (Coolaber, Beijing, China). For inducing sporulation and isolating haploids, *S cerevisiae* transformants were cultured in liquid YPD (1% yeast extract, 2% peptone, 2% glucose) medium for 2 days. Then, the *S. cerevisiae* cells were collected via centrifugation and streaked on McClary medium (0.1 % glucose, 0.18 % KCl, 0.82 % NaAc, 0.25 % yeast extract, 2 % agar) for 7 days at 25°C. To detect the formation of ascospores, cells on the Mcclary medium were stained using Kinyoun and methylene blue on a slide according to the previous method (24). The ascospore was stained red and the thallus of vegetative yeast cells was stained blue under an optical microscope (fig. S1). For haploid isolation, the cells on the McClary medium (ascospore confirmed by staining) were collected and suspended in sterile water. Snailase (A600870, Sangon Biotech, Shanghai, China) was used to digest the cell wall and release the spores. After snailase enzymolysis, the cell suspension was incubated at 58°C to kill the vegetative yeast cells. Finally, the cell suspension was spread on YEP (galactose as the carbon source to induce the expression of the transformed gene) agar plates supplemented with G418 to confirm the presence of the *S. cerevisiae erg10* mutant allele. Colonies were selected randomly and the ploidy was confirmed using PCR (the primers for MATa and MATα are listed in Table S2). The recovered *erg10* mutant was confirmed using PCR with *S. cerevisiae* and *A. oryzae* Erg10/AoErg10s-specific primer pairs. For the temperature-sensitive tests, the control and transformants were grown on YPD (1% yeast extract, 2% peptone, 2% glucose, 1% agar), and YPG (1% yeast extract, 2% peptone, 2% galactose, 1% agar) to assess the phenotypes at 30°C and 37°C. To assess oleic acid consumption, the *pot1* mutant and the corresponding transformants were cultured in the YPD liquid medium supplemented with galactose as the carbon source to induce the expression of the transformed genes. Then, the cells were isolated and inoculated in medium with oleic acid as the sole carbon source.

### Gene Overexpression

All the gene overexpression experiments were performed using the binary vector *pEX2B* (25). To construct the vector, the CDS of *AoErg10s* was fused with *DsRed* at the C-terminal of *AoErg10s* using fusion PCR. The primers used are listed in Supplementary Table S1. The binary vector *pEX2B* was linearized with *Afl*II, followed by cloning of the fusion DNA fragments into *pEX2B* using a one-step cloning kit (Vazyme B0iotech Co., Ltd., China) to construct *pEX2B-AoErg10s-DsRed* vectors. All the constructed vectors were transformed into *A. tumefaciens* AGL1. Then, the vectors were transformed into *A. oryzae* 3.042 △*pyrG* according to the procedure published by our laboratory (19). Spores of single transformants were collected and cultured for further analysis.

### Subcellular Localization Analysis

The constructed pEX2B*-AoErg10s-DsRed* plasmids were transformed into *A. oryzae* 3.042 *△ pyrG* to determine the subcellular localization of the fusion protein. The pEX2B vector with DsRed as the reporter gene was used as the control. To visualize peroxisomes, the SRL amino acid was fused to the C-terminal of GFP to create a peroxisome targeted signal (PTS1) (26), and this GFP sequence was cloned in pEX1-*ptr*A to construct pEX1-*ptr*A-GFP-PTS1 (27). For studying co-localization, the GFP and DsRed vectors were transformed into *A. oryzae* 3.042 *△ pyrG*. For mitochondrial staining, mycelia grown on the slides were transferred into a medium containing Mito-tracker Green (Beyotime Institute of Biotechnology, Nantong, China) and incubated for 30 min at 30°C (28). The mycelia were then washed twice with distilled water and observed using fluorescence microscopy. The fluorescence was observed using a Leica DM4000B microscope. The primer sequences used for plastid construction are shown in Table S2.

### Measurement of Ergosterol and Fatty Acid Contents

Ergosterol and total lipid extraction was performed according to previously described methods (27, 29–31). The 72 h-old *A. oryzae* mycelium was harvested and vacuum freeze-dried to a constant weight. Then, the mycelium was pulverized into powder, and 50 mg dry powder was used for ergosterol extraction and determination according to the following method. The weighted *A. oryzae* powder was suspended in 3 mL alcoholic KOH (25 g KOH plus 35 ml ddH_2_O, with 100% ethanol added to a total volume of 100 mL) by vortexing for 1 min. Then, the suspension was transferred to glass tubes and incubated at 85°C in a water bath for 1.5 h. After the tubes cooled to ambient temperature, 3 mL n-heptane (Sigma-Aldrich, St Louis, MO, USA) and 1 mL distilled water was added to the tube, followed by vigorous vortexing for 3 min to extract ergosterol. The upper layer (n-heptane layer) was isolated and stored at −20°C for 24 h prior to high-performance liquid chromatographic (HPLC) analysis. HPLC was conducted on Waters Alliance e2695-2489 UV/Vis detector HPLC (Milford, MA, USA) using a UV detector set at 282 nm with a Zorbax SB-C18 column. Methanol/water (95:5, v/v) was used as the mobile phase, and the elution rate was 1.5 mL min^-1^. Ergosterol (Sigma-Aldrich) was used to obtain a calibration curve. Each experiment was repeated thrice.

For total lipid extraction and determination, 1 g dry *A. oryzae* powder was placed in a 50 mL Falcon tube. Two milliliters of Triundecanoin (C11:0) (ANPEL Laboratory Technologies (Shanghai) Inc.) internal standard solution (0.5 mg/mL final concentration), 100 mg pyrogallic acid, 2 mL ethyl alcohol, and 4 mL distilled water were added, followed by the addition of two separate zeolites. This was followed by the addition of 10 mL hydrochloric acid solution to the sample, shaking for mixing well, and incubation in a water bath at 75°C for 40 min. The sample was agitated every 5 min to ensure that the solution was homogeneous. After the suspension cooled to room temperature, 10 mL ethyl alcohol was added and mixed well, following which the hydrolyzed blend was transferred to a tube via a separatory funnel. Next, 30 mL 1:1 ether-petroleum ether was added, the tube was covered with a lid and shaken for 5 min, and then allowed to stand for 10 min for the phases to separate. The extraction of the hydrolysate blend was repeated twice. Subsequently, the ether layer extract solution was separated and the lipid extract (organic phase) was dried in a Rotovapor and dissolved in 8 mL of 2% sodium hydroxide methanol solution. The lipid extracts were incubated in reflux condenser at 80 °C for 20 min and 7 mL of 15% boron trifluoride methanol solution was added and incubated at 72 °C for 10 min to obtain fatty acid methyl esters (FAMEs). After the solution cooled to room temperature, the FAME components were extracted using 10 mL n-heptane. A mixed fatty acid methyl ester standard (ANPEL Laboratory Technologies (Shanghai) Inc.) was used to plot a standard curve. Samples were separated and analyzed using a gas chromatographic column (Agilent 7890B GC system) equipped with an Agilent J&W HP-88 GC column (100 m×0.25 mm ID×0.2 μm). The Agilent OpenLAB CDS software was used to extract peak area and retention time. Fatty acid contents were calculated using the standard curve and are expressed as mg/g dry weight cells.

## Results

### 1. Bioinformatics characterization of thiolases in *A. oryzae*

The *S. cerevisiae* thiolase protein (Erg10 and Pot1) sequences were used as queries for BLAST analysis of the corresponding homologous proteins in *A. oryzae* genome in NCBI (http://www.ncbi.nlm.nih.gov/). Six thiolases (named AoErg10A-AoErg10F) were identified; the protein length ranged from 409 to 431 amino acids, with Thiolase_N and Thiolase_C functional domains (table 1) and the functional motifs were predicted (fig. S2). To obtain more information regarding these six *A. oryzae* thiolases, we performed phylogenetic analysis of thiolases in different organisms, including *S. pombe*, *S. cerevisiae*, *B. graminis*, *B.taurus*, *N. tabacum*, *Z. mays*, *A. thaliana*, *G. gallus*, *R. norvegicus*, *H. sapiens*, *B. taurus*, *C. lupus familiaris*, *M. musculus*, *Y. lipolytica*, and *E. caballus*. These thiolases can be divided into four types: cytoplasmic thiolase II (acetyl-CoA C-acetyltransferase), mitochondrial thiolase II (acetyl-CoA C-acetyltransferase), mitochondrial thiolase I (3-ketoacyl-CoA thiolase), and peroxisomal thiolase I (3-ketoacyl-CoA thiolase). As showed in Figure 1, AoErg10A belongs to the branch of cytoplasmic acetyl-CoA C-acetyltransferase, AoErg10B was located at the branch of cytoplasmic and mitochondrial acetyl-CoA C-acetyltransferase, and AoErg1C-AoErg10F was located at the branch of mitochondrial and peroxisomal 3-ketoacyl-CoA thiolase. However, when other thiolases were analyzed, all the thiolases from *A. oryzae* were found to form an independent sub-branch, indicating that the function of these thiolases in *A. oryzae* may differ from those of other species.

**Figure 1.**
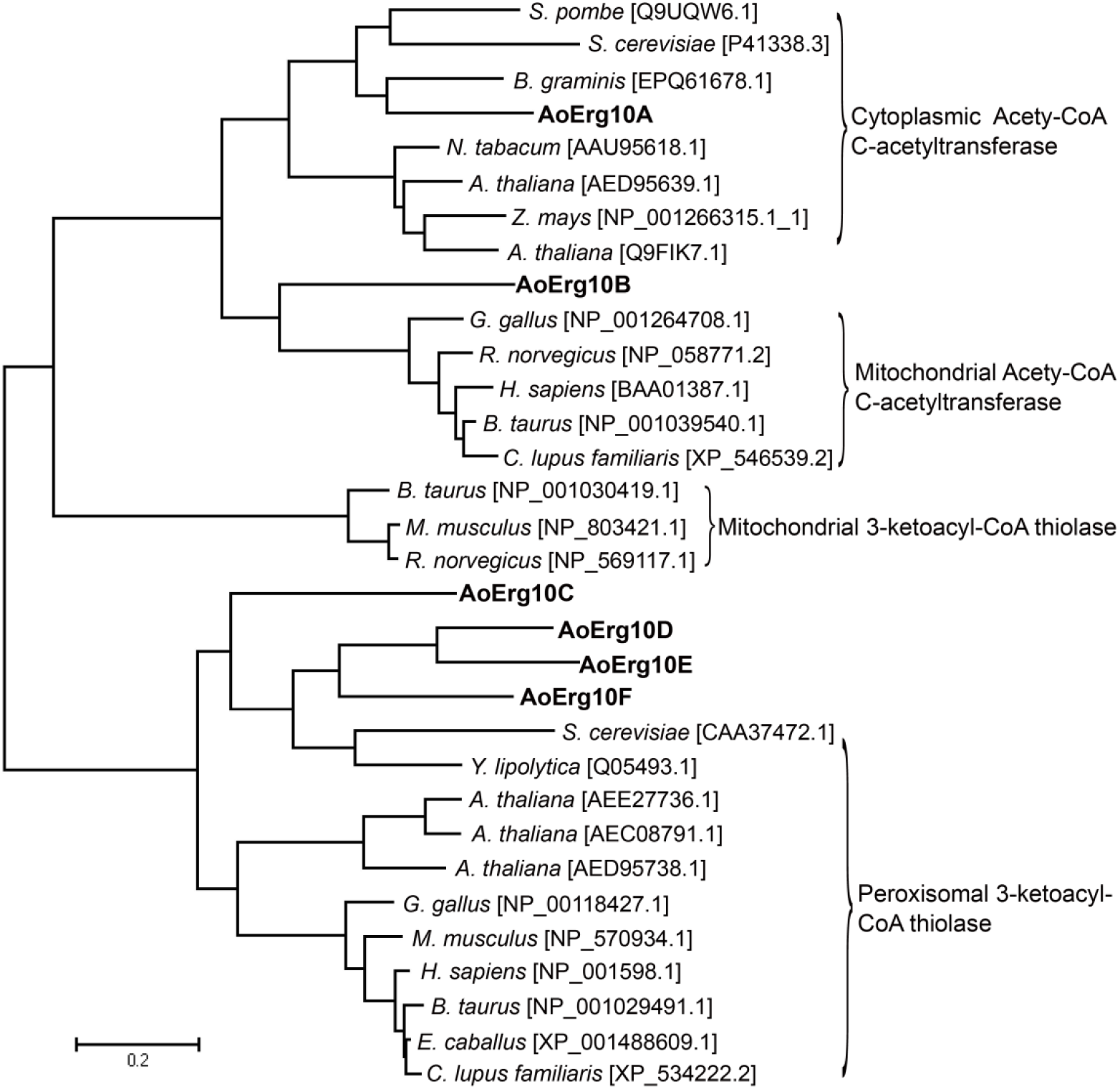
Phylogenetic analysis of thiolase proteins in different species. Unrooted phylogenetic tree of thiolase and homologous proteins in *S. pombe*, *S. cerevisiae*, *B. graminis*, *B.taurus*, *N. tabacum*, *Z. mays*, *A. thaliana*, *G. gallus*, *R. norvegicus*, *H. sapiens*, *B. taurus*, *C. lupus familiaris*, *M. musculus* and *Y. lipolytica*, and *E. caballus*. The IDs of the sequences were included after the specie names in the figure.

**Table 1.**
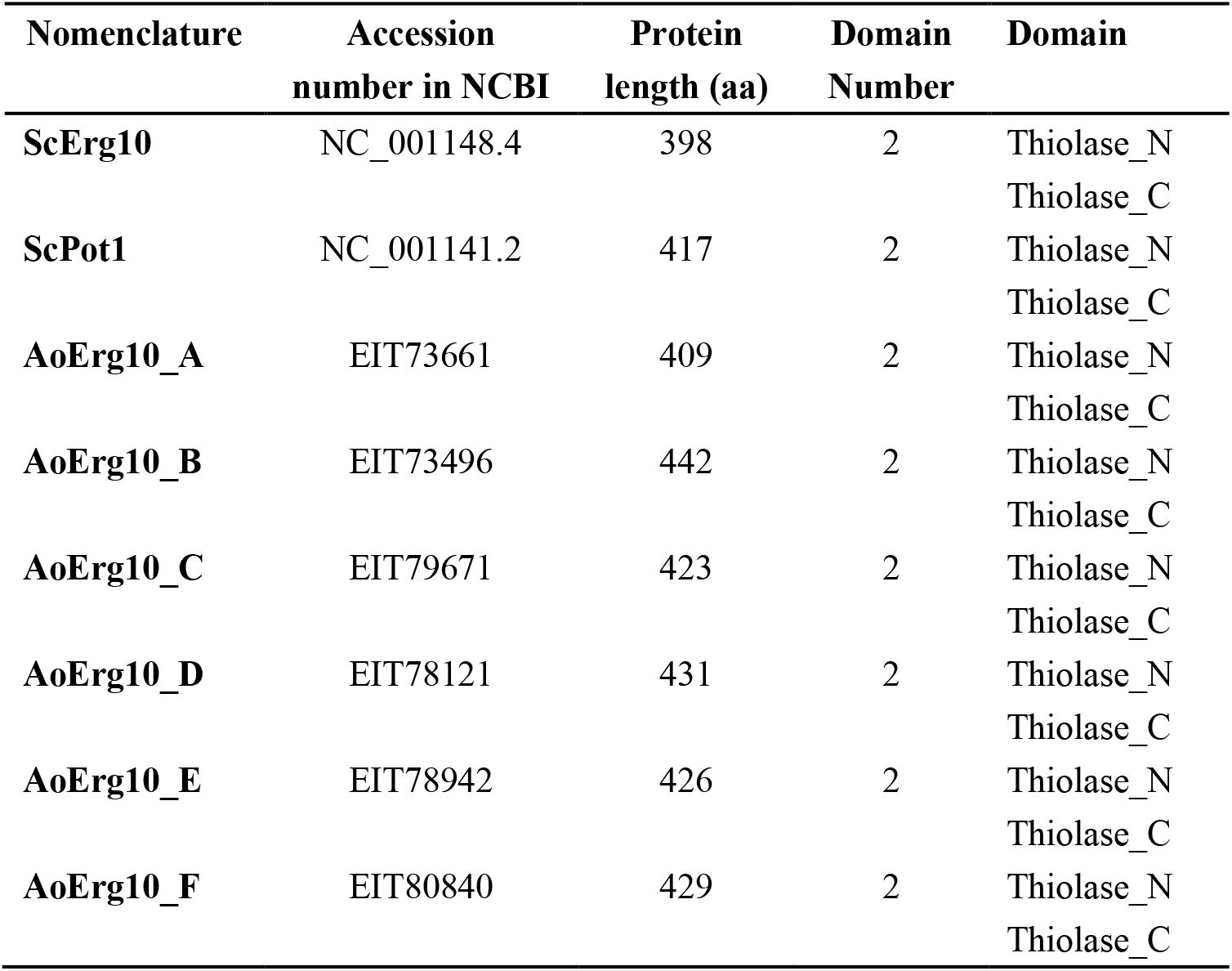
The identification of thiolase proteins in A. oryzae

### 2. Expression pattern of thiolase-encoding genes

To investigate the role of these thiolases during *A. oryzae* growth, we determined the expression patterns of *AoErg10s* (represent for *AoErg10A* to *AoErg10F*) at different developmental stages using qRT-PCR. As shown in Figure 2A, under normal conditions, the expression of these thiolase genes differed at different growth stages. For example, *AoErg10A* is the predominant thiolase at 24 h, which was almost five times higher than that of *AoErg10B* and 10 times more than that of *AoErg10C to AoErg10E*, while *AoErg10F* showed negligible expression. These genes can be divided into two groups based on their expression levels at 48 h: group one included *AoErg10A*, *AoErg10B,* and *AoErg10F,* and showed nearly similar expression, while group two included *AoErg10C*, *AoErg10D*, and *AoErg10E* and showed similar expression; the expression level in group one was nearly five times of that in group two. The expression levels changed at 72 h: the highest expression level was observed for *AoErg10B*, followed by *AoErg10A* and *AoErg10F.* In terms of relative expression level, *AoErg10A* and *AoErg10B* are the predominant thiolases at 24 h, while *AoErg10A*, *AoErg10B*, and *AoErg10F* are mainly expressed after 24 h.

**Figure 2.**
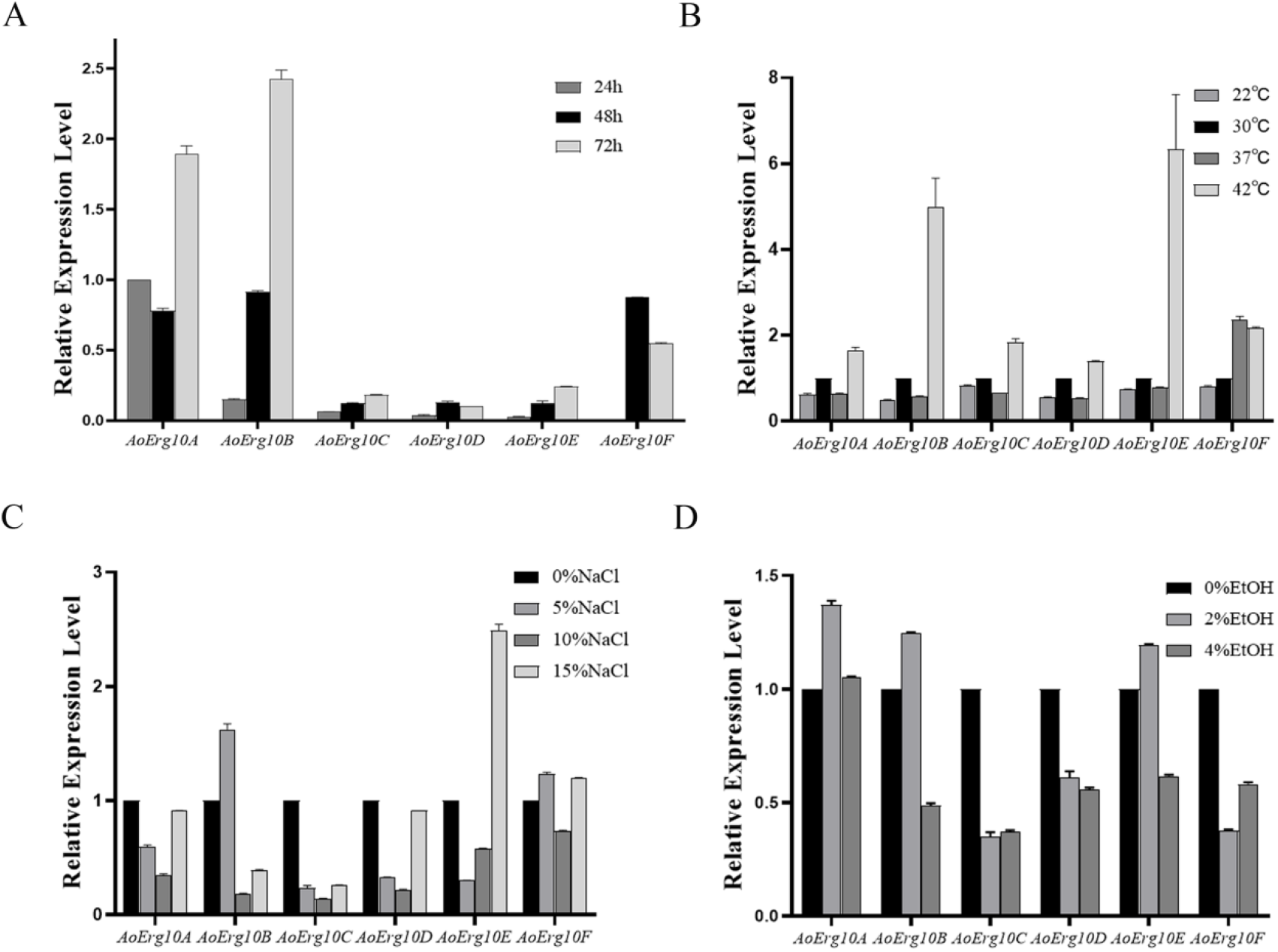
Expression levels of *AoErg10A* to *AoErg10F* at different developmental stages and under different abiotic stress. (A) Expression of *AoErg10A* to *AoErg10F* at 24, 48, and 72 h of growth; (B–D) Expression of *AoErg10A* to *AoErg10F* under temperature, salt and ethanol stress conditions. The wild-type *A. oryzae* spore suspension was plated on CD agar medium or CD agar medium supplemented with NaCl or ethanol and incubated at 30°C (except for temperature stress). For the determination of *AoErg10A* to *AoErg10F* mRNA levels in different growth stages, mycelia were harvested at 24, 48, and 72 h, and the *AoErg10A* expression level at 24 h was used as the reference. For other tests, the mycelia were harvested at 72 h, and the mRNA of the corresponding untreated control or 30°C were used as references. Values represent the mean ± SD of three independent experiments. The primers used are shown in Table S1.

Thiolases are involved in lipid synthesis and metabolism, which are important for stress responses in many organisms (32). Therefore, we also investigated the expression of these thiolase genes under different stress conditions. *A. oryzae* was subjected to temperature, salt, and ethanol stress. For temperature stress, *A. oryzae* was grown at 22°C, 30°C, 37°C, and 42°C. At 22°C, the expression of *AoErg10A*-*F* decreased; at 37°C, the expression levels of *AoErg10A*-*E* decreased, while that of *AoErg10F* increased. However, at 42°C, almost all genes were up-regulated, the most up-regulated being *AoErg10B* and *AoErg10E* (although the expression of *AoErg10E* was low at 30°C) (Fig. 2B). For salt treatment, *A. oryzae* was grown in medium supplemented with 0%, 5%, 10%, and 15% NaCl. The expression levels of *AoErg10A*, *AoErg10C*, *AoErg10D*, and *AoErg10E* in the presence of 5% and 10% NaCl were lower than that without NaCl. *AoErg10B* and *AoErg10F* showed high expression under 5% NaCl stress and low expression under 10% NaCl stress. Under 15% NaCl stress, the expression of *AoErg10A* and *AoErg10D* changed significantly; the expression of *AoErg10B* and *AoErg10C* decreased, whereas that of *AoErg10E* and *AoErg10F* increased (Fig. 2C). For ethanol stress, *A. oryzae* was grown in medium supplemented with 0%, 2%, and 4% ethanol. Results showed that the expression of *AoErg10A*, *AoErg10B,* and *AoErg10D* increased slightly under 2% ethanol stress, while those of the other genes decreased under 2% and 4% ethanol stress (Fig. 2D). Conclusively, *A. oryzae* harbors six thiolase genes, expression levels of which varied at different growth stages and in response to different abiotic stress, indicating the complex distribution of these thiolases.

### 3. Response of thiolase expression to oil acid

Previous studies have shown that the expression of *Pot1* in *S. cerevisiae* can be induced by oil acid and inhibited by glucose (8, 11). Therefore, we investigated the expression levels of these six genes using glucose, glycerol, and oil acid as carbon sources. As showed in Figure 3, the expression of all six genes was slightly inhibited in glucose medium compared to that in medium containing glycerol as the carbon source. *AoErg10B*, *AoErg10D, AoErg10E* and *AoErg10F* were significantly induced on oil acid medium, while the expression of *AoErg10D*, *AoErg10E*, and *AoErg10F* increased by nearly 5-6 times compared to that on glycerol medium. The expression of *AoErg10E* was about 3 times higher than that on glycerol medium. Thus, among these six genes, *AoErg10B, AoErg10D*, *AoErg10E,* and *AoErg10F* showed similar response to oil acid as *Pot1* in *S. cerevisiae*, indicating that these genes may be involved in β-oxidation of fatty acids.

**Figure 3.**
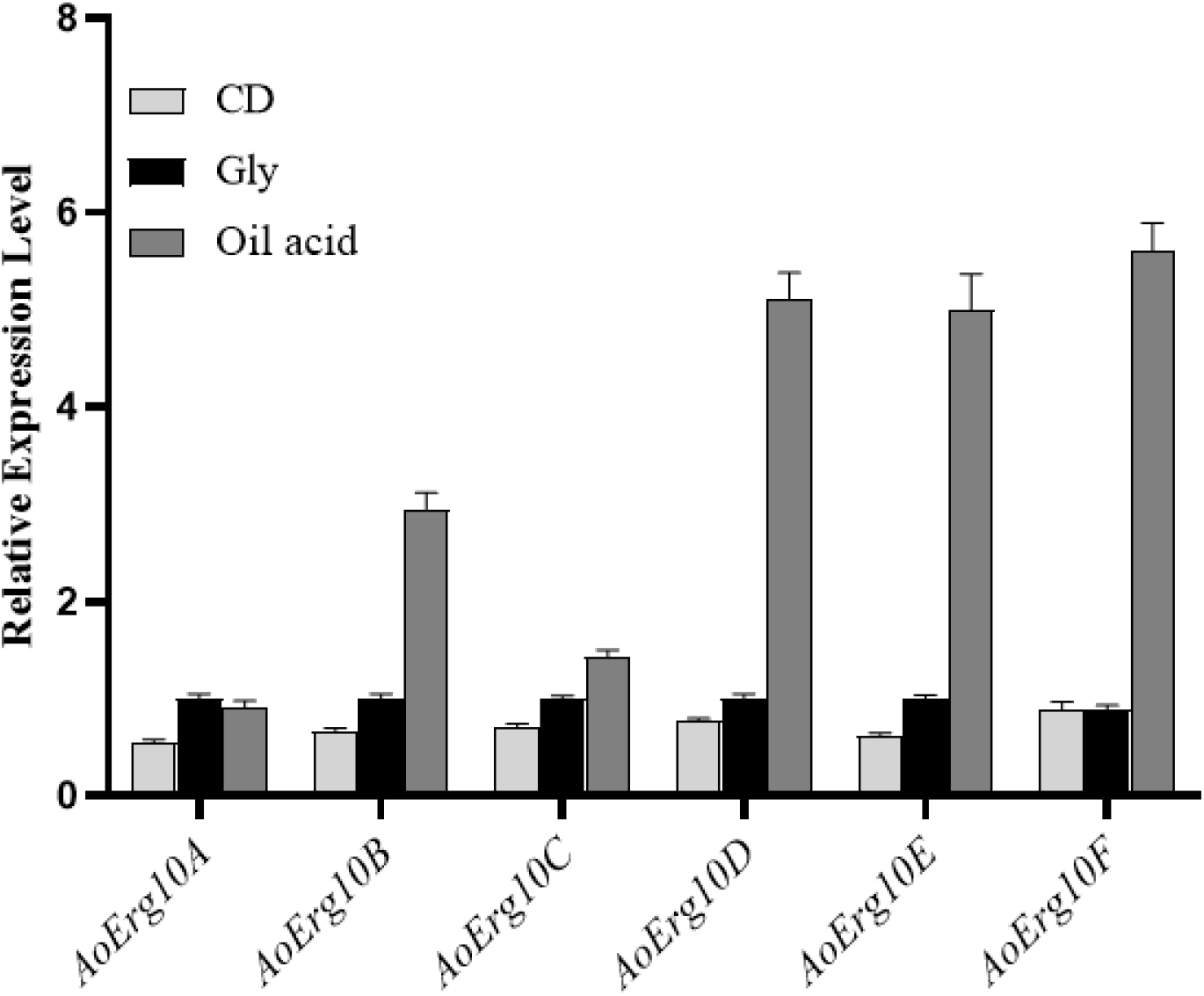
Expression of *AoErg10s* after glycerol and oleic acid treatment. *A. oryzae* was cultured on the CD medium with cellophane for 24 h and transferred to medium with glycerol or oleic acid as carbon source and incubated for 10 h. Then, the mycelia were harvested for RNA isolation and gene expression analysis. The mRNA levels of *AoErg10s* in CD medium were used as references.

### 4. Subcellular localization of six thiolases

Identification of the subcellular localization of the six thiolases is critical for understanding their function. Although theses thiolases were phylogenetically divided into cytoplasmic-, mitochondrial- and peroxisomal-localized thiolases, the results of subcellular localization analysis showed negligible differences. For example, mitochondria-targeting amino acid sequences (mts) are present at the N-terminus of AoErg10B and AoErg10C, while AoErg10C was categorized into peroxisomal-localized thiolases based on phylogenetic analysis. In addition, specific targeting signal sequences were absent in the amino acid sequence of the other AoErg10s. Hence, we used GFP or DsRed as the reporter gene to investigate the subcellular localization of AoErg10s in *A. oryzae*. The DsRed gene was fused at the C-terminus of AoErg10s to construct the overexpression vectors using uridine/uracil auxotrophic as the selection marker. Meanwhile, the peroxisome (PTS-GFP)- and mitochondria (MTS-GFP)-targeted GFP vectors using pyrithiamine as selection markers were also constructed as reported previously (27). We observed that AoErg10A was distributed uniformly in the cell, indicating that the cytosolic localization of AoErg10A; AoErg10B-DsRed and AoErg10C-DsRed showed irregular and mobile structures reminiscent of the mitochondria; the fluorescence of AoErg10D-DsRed– AoErg10F-DsRed showed punctate structure, which was similar to that of peroxisomes (Fig. 4A).

**Figure 4.**
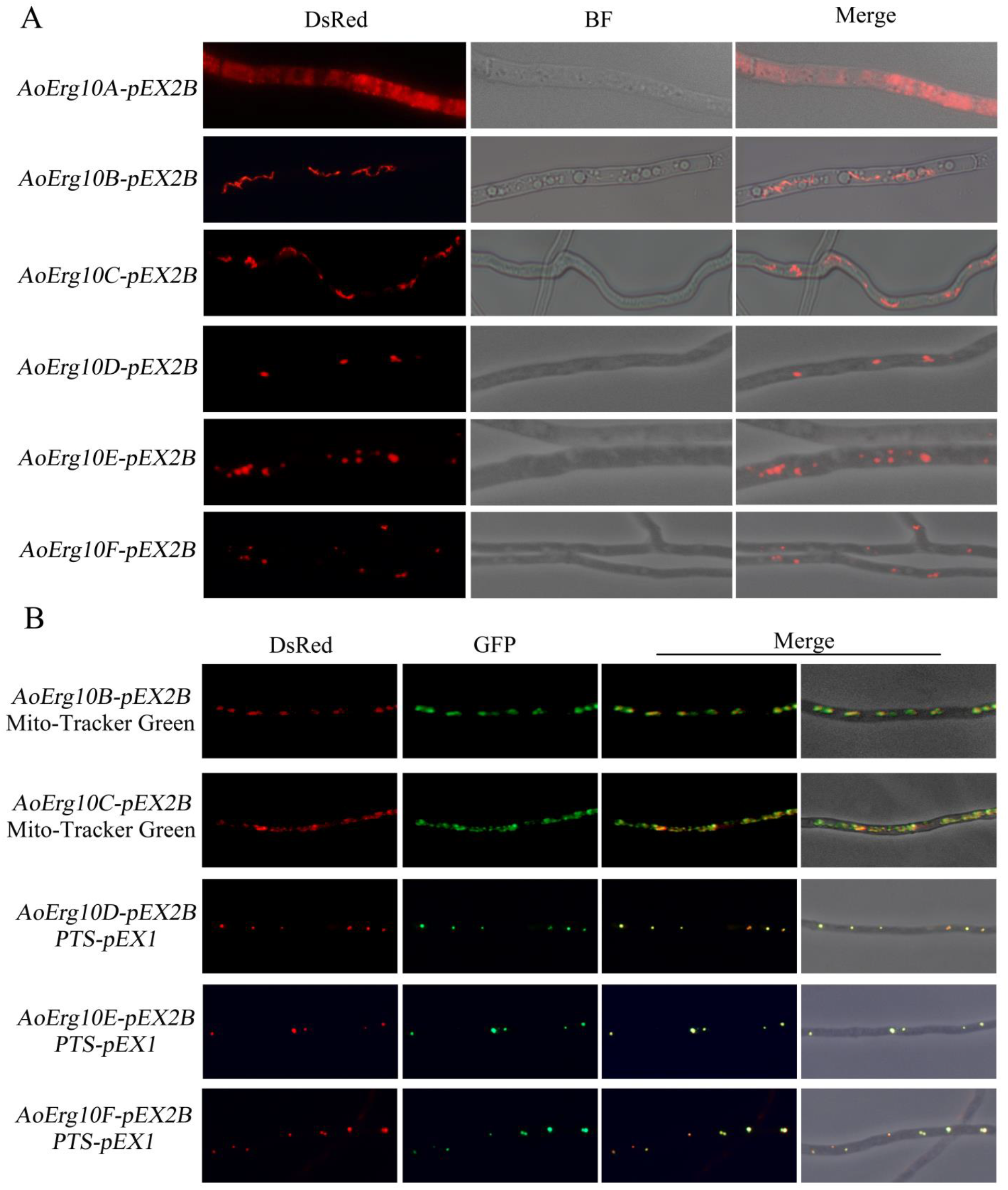
Subcellular localization of AoErg10s. (A) The mycelium of *A. oryzae* 3.042 *△pyrG* transformed with AoErg10s-DsRed. Left to right: fluorescent image of DsRed, bright field, and merged image of DsRed and bright field. (B) Co-localization of AoErg10B-DsRed-AoErg10F-DsRed with peroxisome and mitochondria. The mycelium of *A. oryzae* 3.042 *△pyrG* transformed with AoErg10B-DsRed and AoErg10C-DsRed strains are stained with the mitochondrial marker dye, MitoTracker-green. The mycelium of *A. oryzae* 3.042 *△pyrG* were co-transformed with AoErg10D-DsRed-AoErg10F-DsRed and PTS-GFP vectors. Left to right: fluorescent image of DsRed, GFP, merged image of DsRed and GFP, and merged image of DsRed, GFP and/or bright field.

Consistently, the AoErg10B-DsRed and AoErg10C-DsRed strains were stained with the mitochondrial marker dye, MitoTracker-green, and showed co-localization of red and green fluorescence, indicating that AoErg10B and AoErg10C localized to the mitochondria (Fig. 4B). The AoErg10D-DsRed–AoErg10F-DsRed strains were co-transformed with peroxisome-targeted GFP vectors. The results showed that the red and green fluorescence signals co-localized in all AoErg10D–AoErg10F transgenic strains, indicating that these three thiolases were localized in the peroxisome (Fig. 4B). We also observed that the AoErg10B-DsRed and AoErg10C-DsRed did not co-localize with peroxisome-targeted GFP fluorescence (Fig. S3A). In addition, we deleted the mts of AoErg10B and AoErg10C to construct *AoErg10B*^ΔMTS^-DsRed and *AoErg10C*^ΔMTS^-DsRed vectors. Microscopy showed that the red fluorescence signals were uniformly distributed in the cytoplasm (Fig. S3B). Therefore, we concluded that AoErg10A is a cytosolic thiolase; AoErg10B and AoErg10C targeted the mitochondria due to their mitochondria-targeted amino acid sequences, and AoErg10D–AoErg10F were peroxisomal thiolases.

### 5. AoErg10s recovered the phenotype of the *S. cerevisiae* thiolase II mutant

Yeast heterologous complementation assay was performed to identify whether these thiolases functioned similar to Erg10, the only thiolase II in *S. cerevisiae*. *Erg10* is an essential gene and a homozygous loss of function mutant is lethal. Hence, a weak *S. cerevisiae erg10* (Y40985) mutant that showed temperature-sensitive phenotype was used for the yeast heterologous complementary assay. Full length CDSs of the six genes were cloned into the yeast expression vector, pYES2.0, and transformed into the *erg10* weak mutant. As pYES2 contains a galactose-induced *GAL1* promoter, all the transformants were grown on YPD (with glucose) and YPG (with galactose as an inducer) media at 30°C and 37°C to observe the phenotype. It is interesting that not only AoErg10A, but also AoErg10D, AoErg10E, and AoErg10F can restore the lethal phenotype of the *erg10* weak mutant at 37°C (Fig. 5A). As AoErg10B and AoErg10C were located in the mitochondria, we deleted the mts of AoErg10B and AoErg10C to construct *pYES2.0-AoErg10B^ΔMTS^* and *pYES2.0-AoErg10C^ΔMTS^* vectors and transformed them in the *erg10* weak mutant. The results showed that only *AoErg10B^△mts^* can restore the temperature-sensitive phenotype (Fig. 5B). As evolutionarily AoErg10C is close to peroxisome-targeted thiolase I (Fig. 1), we also add the PTS sequences at the C-terminal of AoErg10C (with or without the mts); results showed that neither AoErg10C^ΔMTS^+PTS nor AoErg10C+PTS can restore the temperature-sensitive phenotype of the *S. cerevisiae erg10* weak mutant (Fig. 5B). The growth curve at 37°C in YPG liquid medium was also plotted. Consistent with the growth observed in solid medium, AoErg10A, AoErg10B*^ΔMTS^*, AoErg10D, AoErg10E, and AoErg10F recovered the temperature-sensitive phenotype of the *S. cerevisiae erg10* weak mutant. Although their growth rates differed, all the recovered transformants reached the same optical density at stationary phase (Figure 5C). Furthermore, we also measured the ergosterol content in all transformants and found that compared to that of the mutant, the ergosterol content increased in almost of the transformants to different levels; however, their ergosterol content was still lower than that of the wild type. Among these transformants, the ergosterol content of *AoErg10B*/*erg10*, *AoErg10C*/*erg10* and *AoErg10C^ΔMTS^*/*erg10* were lower than those of others, which is consistent with their phenotypes at 37°C (Figure 5D).

**Figure 5.**
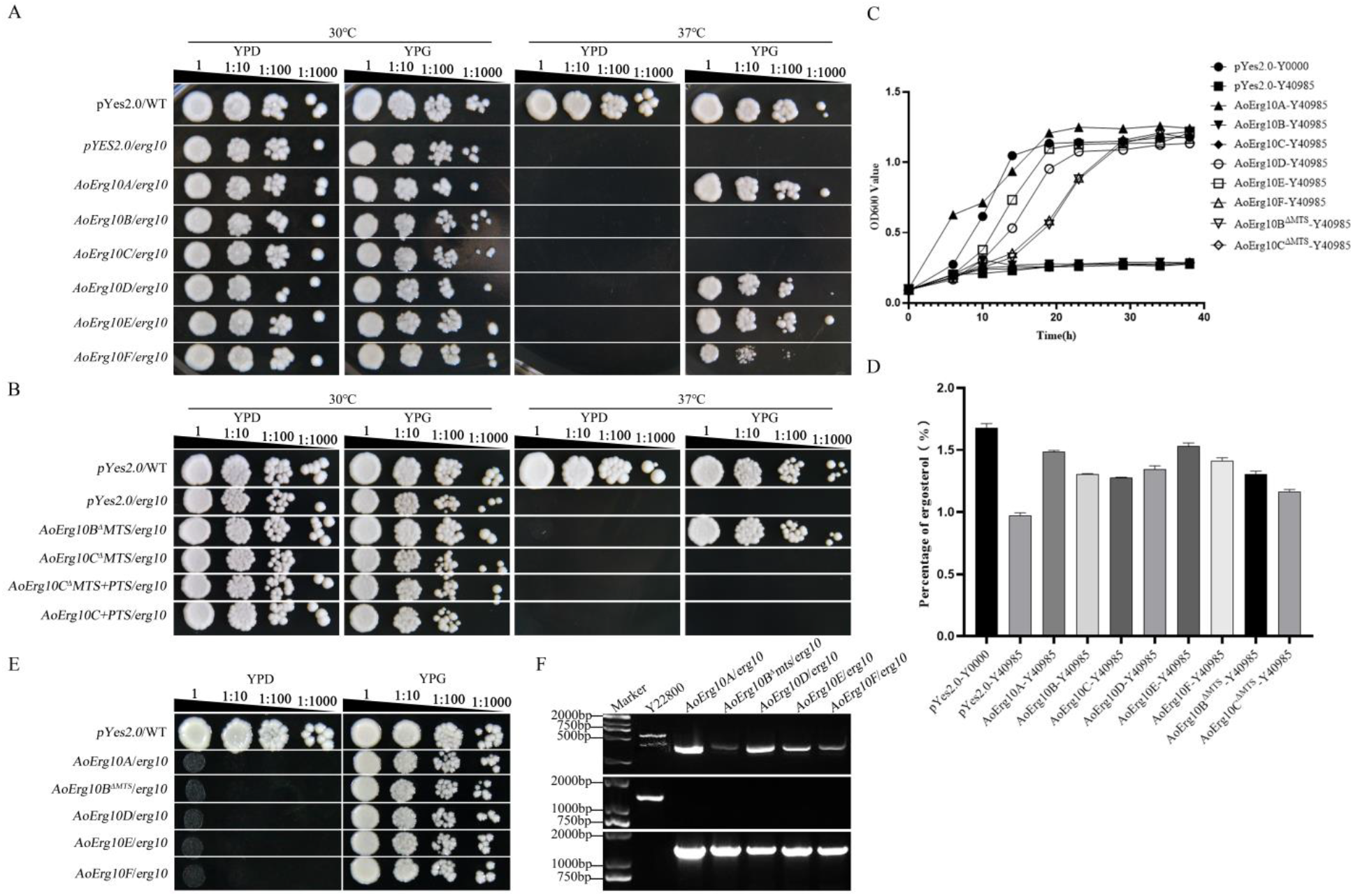
*AoErg10s* recovered the phenotypes of the *erg10* mutant of *S. cerevisiae*. (A) Growth of wild type, *erg10* weak mutant (Y40985) and *AoErg10s*/*erg10* transformants on YPD and YPG medium at 30°C and 37°C. (B) Growth of wild type, *erg10* weak mutant (Y40985), and A*oErg10B*/*C* with different subcellular localization transformants on YPD and YPG medium at 30°C and 37°C. (C) Growth curves of wild type, *erg10* weak mutant (Y40985), and *AoErg10s* in YPG medium at 37°C. (D) Ergosterol content in wild type, *erg10* weak mutant (Y40985), and *AoErg10s* transformants. (E) The phenotypes of isolated haploid *S. cerevisiae erg10* lethal mutant. (F) PCR confirmation of isolated haploid transformants with *erg10* deficiency. The constructs of A*oErg10A*, *AoErg10B^ΔMTS^*, *AoErg10D*, *AoErg10E*, and *AoErg10F* were transformed into heterozygote diploids with copy of loss of function *Erg10*. Then, haploid transformants with *Erg10* loss of function were isolated and grown on YPD or YPG medium. The ploidy was confirmed using PCR primers for MATa and MATα. *Erg10* was confirmed using specific primers and the transformed genes were confirmed using *AoErg10s* specific primers (Table S2).

Phylogenetic analysis showed that AoErg10D, AoErg10E, and AoErg10F belong to type I peroxisomal thiolase. Two possible mechanisms can explain the complementation of the *erg10* weak mutant. First, the transgenic type I peroxisomal thiolase enhanced the fatty acid β-oxidation and increased the concentration of acetyl coenzyme A (the substrate of Erg10), increasing the biosynthesis of ergosterol. Second, phylogenetic peroxisomal thiolase I (AoErg10D, AoErg10E, and AoErg10F) can catalyze the condensation of two molecules of acetyl-CoA to form acetoacetyl-CoA, similar to Erg10 in *S. cerevisiae*. To validate this hypothesis, we used another *erg10 S. cerevisiae* mutant (Y22800), which is a heterozygous diploid yeast cell containing a copy of loss of function *Erg10,* as the homozygous mutant was lethal. We transformed the vectors (A*oErg10A*, *AoErg10B^ΔMTS^*, *AoErg10D*, *AoErg10E,* and *AoErg10F*) that can restore the temperature-sensitive phenotype of the weak mutant into the heterozygous diploid yeast cell. Haploid transformants with *Erg10* loss of function can be isolated from the transgenic heterozygous diploid if these genes can function similar to Erg10 in yeast. Our results showed that the *erg10* loss of function haploid cells can be isolated from the heterozygous diploid transformants, and that the transformed *erg10* loss of function haploid can only grow in medium after induction of *AoErg10s* expression by galactose (Fig. 5E and 5F). To test the functional specificity of the thiolases in *S. cerevisiae*, the known type I thiolase gene, *Pot1* was used to complement the *erg10* weak mutant. We found that *Pot1* cannot restore the temperature-sensitive phenotype of the *erg10* mutant (Fig. S4). Therefore, these results indicated that not only phylogenetic thiolase II (AoErg10A and cytosolic AoErg10B^ΔMTS^), but also phylogenetic peroxisomal thiolase I (AoErg10D, AoErg10E, and AoErg10F) can act as cytosolic thiolase II to recover ergosterol biosynthesis in *S. cerevisiae*.

### 6. AoErg10s recovered the phenotypes of *S. cerevisiae* thiolase I mutant

We next identified whether these *AoErg10s* can function as thiolase I to catalyze the last step of fatty acid β-oxidation. The *S. cerevisiae* thiolase I mutant, *pot1* was used for heterologous complementation assay. Pot1 is the only thiolase I in *S. cerevisiae*, and the *pot1* mutant cannot survive in medium with oleic acid as the carbon source (11). All *AoErg10s* were transformed into *pot1* to test whether these genes can catalyze fatty acid β-oxidation. The *AoErg10s/pot1* transformants were cultured in YNB (with 0.1% galactose to induce the transformed gene) supplemented with oleic acid as the only carbon source. The results showed that *AoErg10D*/*pot1* grew well in medium with oleic acid as the only carbon source, and that the concentration of *AoErg10D*/*pot1* was five times that of wild type *S. cerevisiae* (Fig. 6 A and B); the *AoErg10E-* and *AoErg10F-* transformed *pot1* can also grow in medium with oleic acid as the only carbon source, although the concentration was low (Fig. 6 A and B), while the other transformants showed negligible growth, which is similar to that of the *pot1* mutant. As oleic acid floats on the surface of medium, the consumption of oleic acid can be simply observed by the amount of oil droplets. Results showed that large oil droplets can be seen on the surface of the medium of all transformants, with the exception of *AoErg10D*/*pot1*. Thus, among these six thiolases, *AoErg10D*-*F* can function as *Pot1* to metabolize oil acid, similar to that in *S. cerevisiae*.

**Figure 6.**
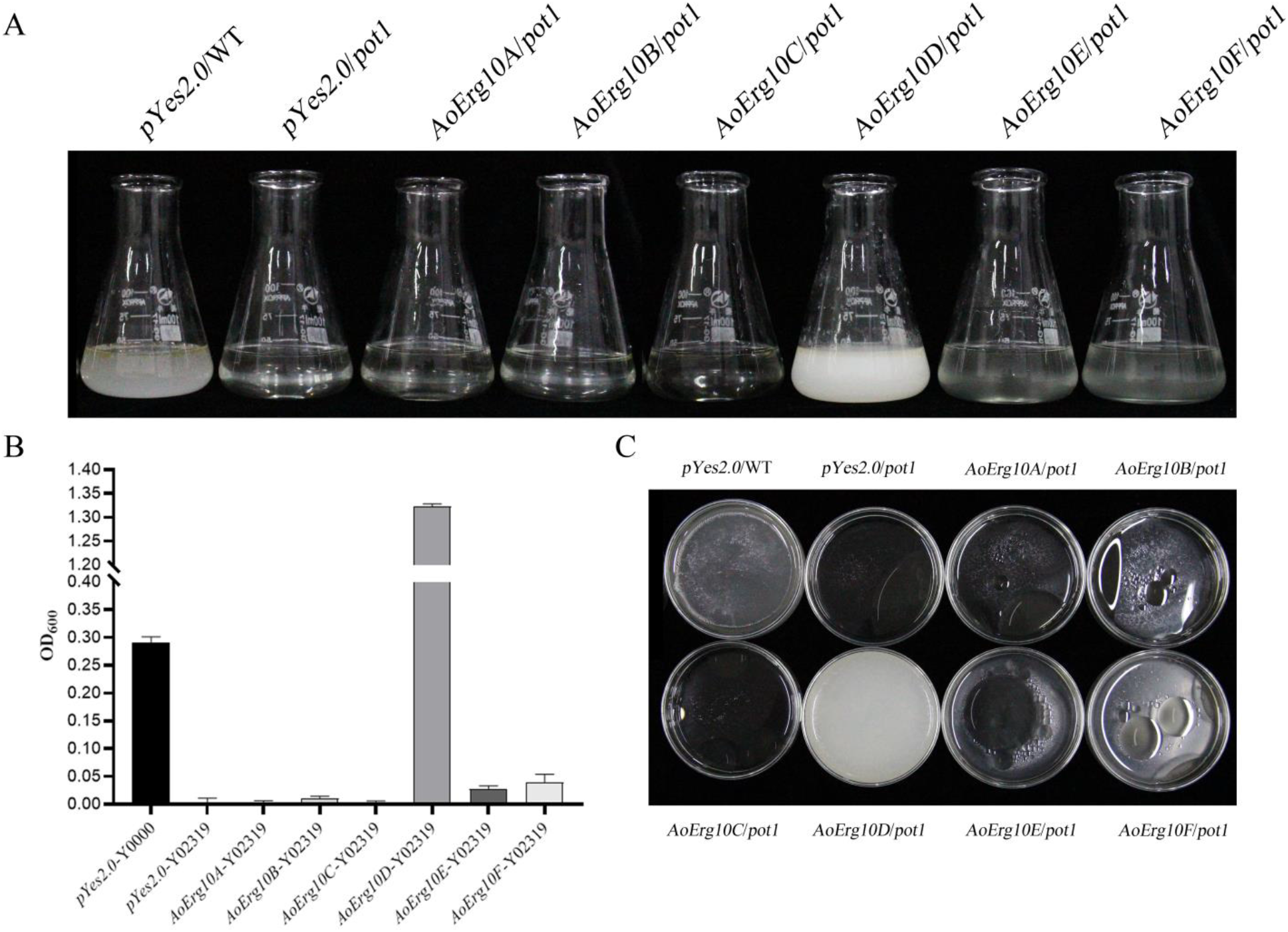
AoErg10s recovered the phenotypes of the *pot1* mutant of *S. cerevisiae*. (A) Growth of wild type, *pYes2.0*/*pot1*, and *AoErg10s*/*pot1* for 72 h in liquid medium with oleic acid as the carbon source. To induce the expression of transformed genes, 0.1% galactose was added to the medium. (B) Liquid medium of wild type, *pYes2.0*/*pot1* and *AoErg10s*/*pot1* after 72 h cultivation. (C) The OD600 of wild type, *pYes2.0*/*pot1*, and *AoErg10s*/*pot1* after 72 h in liquid medium with oleic acid as the carbon source.

### 7. Phenotypes of AoErg10s overexpression strains

All the *AoErg10s* constructs (including *AoErg10B*^ΔMTS^-DsRed and *AoErg10C*^ΔMTS^-DsRed) were transformed into *A. oryze* to obtain the overexpression strains, which were cultured in PDA, DPY, and CD medium (Fig. 7 and S5). The colony morphologies of these strains did not differ remarkably. However, spore number and colony diameter differed. We counted the spore number and colony diameter in PDA medium. As shown in Figure 7, the spore number of *AoErg10A* and *AoErg10C*^ΔMTS^ over expressionstrains decreased, and that of *AoErg10B* increased.

**Figure 7.**
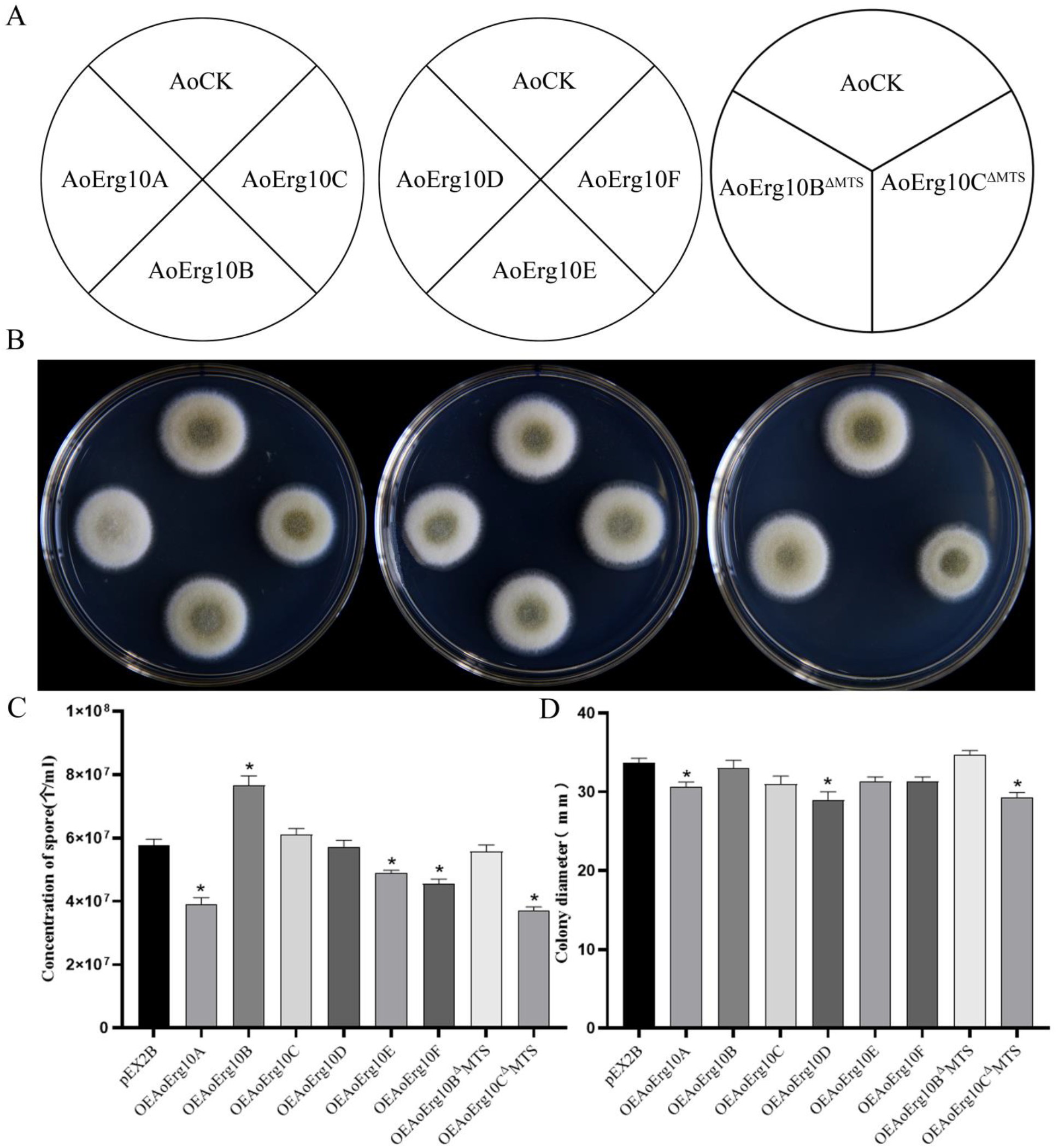
Phenotypes of *AoErg10s* overexpression strains. (A) Scheme showing different transgenic strains. (B) Colony morphologies of the control (AoCK, wild type *A. oryzae*) and AoErg10s overexpression strains on PDA medium after 48 h of incubation. Spore suspension of identical concentrations of different *A. oryzae* strains were plated on PDA agar medium and incubated at 30°C for 72 h. (C and D) Spore number and colony diameter of different transgenic strain colonies.

The colony diameter of *AoErg10A*, *AoErg10D,* and *AoErg10C*^ΔMTS^ overexpression strains decreased, while only the colony diameter of the *AoErg10B*^ΔMTS^ overexpression strain increased. Therefore, we concluded that overexpression of AoErg10s either affected the growth rate or sporulation of the transgenic strains.

### 8. Fatty acid and ergosterol contents of AoErg10s overexpression strains

As thiolase is involved in fatty acid or ergosterol metabolism, we measured the fatty acid and ergosterol contents of all the AoErg10s transgenic strains. As shown in Table 2, the fatty acid content changed significantly in these strains. For example, in *AoErg10A*, *AoErg10D,* and *AoErg10C*^ΔMTS^ overexpression strains, the total fatty acid content decreased to 67%, 84%, and 94% of the control, respectively, while those in the others increased from 3% to 18% compared to that of the control. The unsaturated fatty acid (UFA) and saturated fatty acid (FA) contents also changed in most of these strains. In addition, the composition of fatty acids also changed. Certain fatty acids in CK cannot be detected in the overexpression strains. For example, the C4:0 fatty acid was only detected in CK, *AoErg10A,* and *AoErg10B* overexpression strains, and the concentration in *AoErg10B* overexpression strain was 40 times that in CK (table 1); C17:1 and C22:6 were detected in all strains except for in the *AoErg10B* overexpression strain; C24:0 was detected in CK, *AoErg10C*, *AoErg10F*, *AoErg10B*^ΔMTS^, and *AoErg10C*^ΔMTS^ overexpression strains, but not in the other strains. In contrast, the concentrations of composite fatty acids changed. For example, concentration of C22:2 decreased to about half of that in CK in *AoErg10A* and *AoErg10D* overexpression strains. In addition, we also estimated the ergosterol content in these transgenic strains. Compared to that in CK, the ergosterol content in *AoErg10C*, *AoErg10F* and *AoErg10B^ΔMTS^* overexpression strains increased by 11%, 16%, and 8.7%, respectively, while that in AoErg10D, AoErg10E, and AoErg10C^ΔMTS^ decreased by 13%, 15%, and 20%, respectively. Taken together, overexpression of *AoErg10s* disrupted the balance between fatty acid and ergosterol biosynthesis, which affected their cellular content.

**Table 2.**
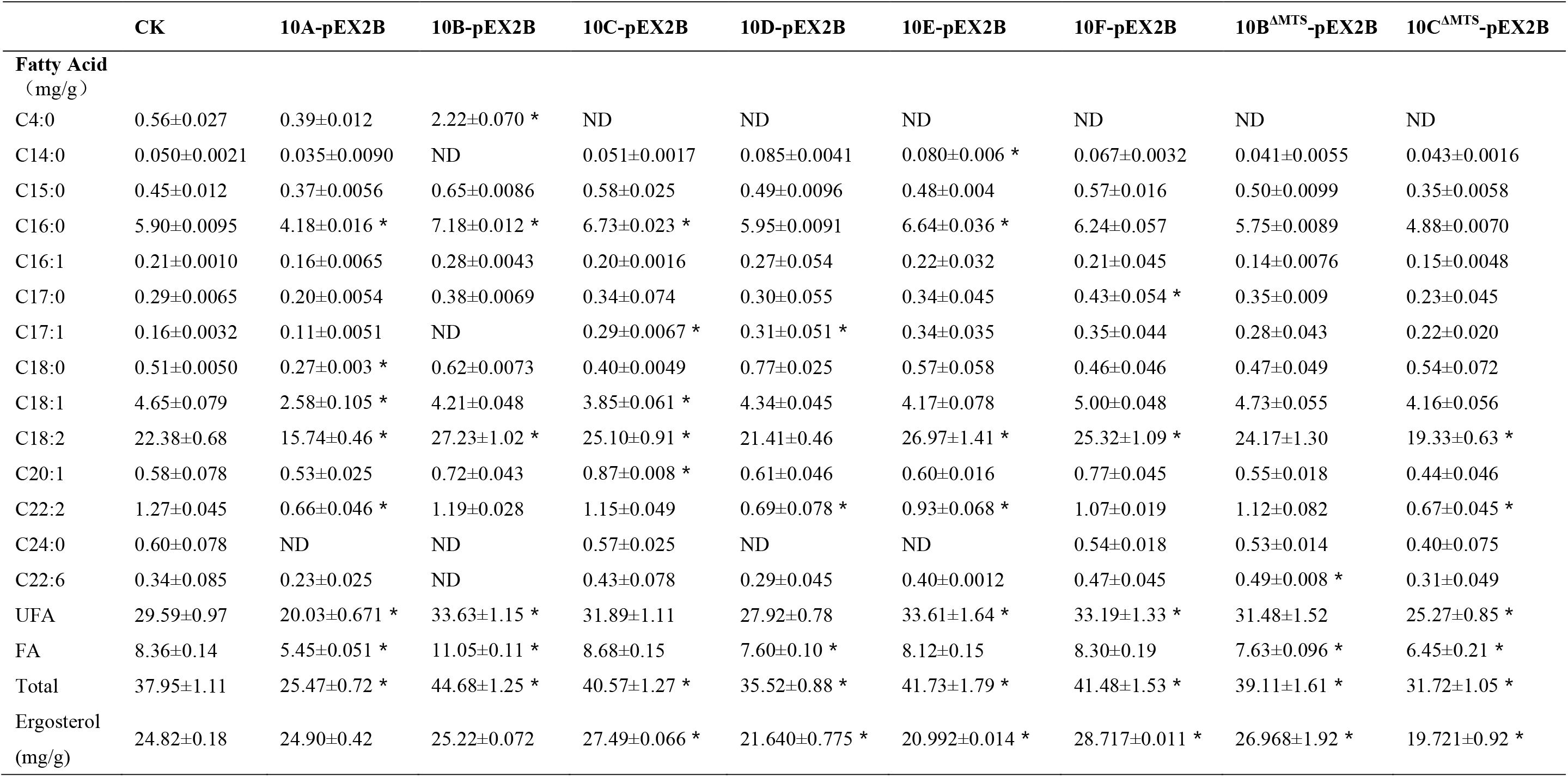
The fatty acid and ergosterol contents in AoErg10 over-expression strians

## Discussion

Thiolase II catalyzes carbon–carbon bond formation via a thioester-dependent Claisen condensation reaction mechanism, which is an essential step in fatty acid and polyketide biosynthesis, while thiolase I catalyzes β-oxidation of fatty acids, which is essential for acetyl-coA and energy production (33). Previous studies have shown that the two classes of thiolases have related sequences and essentially use the same active site residues to perform the relevant reactions, implying their origin from a common ancestor (2). In most situations, the two classes of thiolase can catalyze both degradative and biosynthetic reactions; however, the biosynthetic reaction is thermodynamically unfavorable (1). Degradative and biosynthetic enzymes share the same catalytic mechanism in which the catalytic site is formed by four loops, each harboring conserved residues at the active site. In this study, six thiolase (named AoErg10A-AoErg10F) were identified by bioinformatics analysis, which can be divided into cytoplasmic acety-CoA C-acetyltransferase, mitochondrial acety-CoA C-acetyltransferase and peroxisomal 3-ketoacyl-CoA thiolase by phylogenetic analysis. We detected the subcellular localization and expression pattern of these thiolases and thiolase encoded genes. Yeast heterologous complementary revealed the phylogenetic 3-ketoacyl-CoA thiolase can function as cytoplasmic acety-CoA C-acetyltransferase in yeast. Finally, we also detected the effect of over expression these thiolases on the lipid metabolism.

### The dual function and subcellular localization of *A. oryzae* thiolases

On the basis of function and subcellular localization, known thiolases can be divided into cytoplasmic/mitochondrial acety-CoA C-acetyltransferase and mitochondrial/peroxisomal 3-ketoacyl-CoA thiolase. A previous study has revealed that the two classes of thiolase can catalyze both degradative and biosynthetic reactions in most situations; however, the biosynthetic reaction is thermodynamically unfavorable (1). In this study, six thiolases in *A. oryzae* genome were identified using bioinformatics analysis. Phylogenetic analysis showed that these thiolases belong to three branches: cytoplasmic acety-CoA C-acetyltransferase (AoErg10A), mitochondrial acety-CoA C-acetyltransferase (AoErg10B), and peroxisomal 3-ketoacyl-CoA thiolase (AoErg10C-AoErg10F). However, our subcellular localization analysis showed that AoErg10C was localized in the mitochondria, which was different from the result of phylogenetic analysis. Use of weak and loss of function *S. cerevisiae erg10* mutants revealed that both the cytoplasmic and peroxisomal (except the mitochondrial) thiolases can function as acety-CoA C-acetyltransferase (Erg10) in yeast. Deletion of mitochondria targeting amino acid sequences (mts) at the N-terminal of AoErg10B and AoErg10C showed that only AoErg10B can complement the phenotype of the *S. cerevisiae erg10* mutant, indicating that only AoErg10C is a specific 3-ketoacyl-CoA thiolase. The *S. cerevisiae* thiolase I–*pot1* mutant revealed that only AoErg10D-F can function as thiolase I to recover the oleic acid non-utilizing phenotype of *pot1*. Therefore, AoErg10D-F showed dual function of thioase I and thiolase II in *S. cerevisiae*.

### The effect of thiolase overexpression on *A. oryzae* lipid metabolism

In this study, we showed that the subcellular localization of AoErg10B and AoErg10C can affect their function. Thus, the fatty acid and ergosterol contents in all AoErg10s overexpression strains (including AoErg10B^ΔMTS^ and AoErg10C^ΔMTS^) were determined. Indeed, the contents of fatty acid and ergosterol changed in most thiolase overexpressing strains. Maximum decrease and increase in fatty acid contents were observed in AoErg10A and AoErg10B overexpression strains, respectively; while the most decrease and increased inergosterol contents were observed in AoErg10C^ΔMTS^ and AoErg10F overexpression strains, respectively. Interestingly, the ergosterol content decreased in AoErg10C^ΔMTS^ overexpression strains, while it increased in AoErg10C overexpression strains, which further proved that subcellular localization can affect the function of these thiolases in lipid metabolism. The decrease in fatty acid levels in AoErg10A overexpression strains may be because overexpression AoErg10A induces the synthesis of acetoacetyl-CoA from more acety-CoA. However, the ergosterol content in AoErg10A overexpression strain did not increase. This may because the ergosterol pathway involves many enzymes and steps, and other rate-limiting enzymes may regulate ergosterol content. In contrast, the changes in fatty acid and ergosterol also uncovered that complex regulatory mechanism in lipid metabolism in *A. oryzae*. This study identfied six thiolases in *A. oryzae* and revealed their different functions. Unlike other reported thiolases in fungi, three of the six thiolases showed dual function of thioase I and thiolase II in *S. cerevisiae*, indicating the lipid metabolism is more complex in *A. oryzae*. Indeed, prevoius studies showed that *A. oryzae* has a high capability in production of high lipid content and has been used for lipid production (34–38). Thus, the reveal of founction of these thiolases in *A. oryzae* can lay the foundation for genetic engineering for lipid metabolism in *A. oryzae* or other fungi. Other experiments such as gene knock out or enzyme activity assay should be performed to further reveal the function of these thiolases and mechanism via which they regulate lipid metabolism.

## Acknowledgments

We are grateful to Professor Van-Tuan Tran (VNU University of Science, Hanoi, Vietnam) for This study was supported by Natural Science Foundation of Jiangxi Province (20192ACBL20012 and 20212BAB205001), National Natural Science Foundation of China (NSFC Grant NO.: 31700068), Science and Technology Research Project of Jiangxi Provincial Department of Education (Grant No.: GJJ201141), and the youth talent support program of Jiangxi Science & Technology Normal University (2019QNBJRC004).

## Supplementart materials

Table S1 Primers used for qRT–PCR

Table S2 Primers used for vector construction

Fig.S1. Diploid Spore Staining of *Saccharomyces cerevisiae*

Figure S2. Prediction functional motifs of thiolases in *S. cerevisiae* and *A. oryzae*

Figure S3. Subcellular localization of AoErg10 B/C and AoErg10B/C^ΔMTS^

Figure S4. The phenotype of *Pot1/erg10* transfromed *S. cerevisiae*

Figure S5. Growth phenotype of *AoErg10s* over expression strains on DPY and CD medium

## Legends of supplemental figures

Figure S1. Diploid Spore Staining of *Saccharomyces cerevisiae*. After the diploid yeast was sporulated on Mcclary medium for 7 days, strians were stained by Kinyoun and methylene blue. Blue color is the vegetative yeast cell, and the ascospore is stained with red color.

Figure S2. Prediction functional motifs of thiolases in *S. cerevisiae* and *A. oryzae*. Conserved motifs of the thiolase proteins in *saccharomyces cerevisiae* (ScErg10 and ScPot1) and *A. oryzae* (AoErg10A-F). All conserved motifs of the thiolase proteins were identified by the MEME program. Protein sequences are indicated by thin black line, and the conserved motifs are represented by different colored boxes. The length (the number of amino acids) of the protein and motif can be estimated using the scale bar at the bottom.

Figure S3. Subcellular localization of AoErg10B/C and AoErg10B/C^ΔMTS^. (A) The mycelium of *A. oryzae* 3.042 *△pyrG* were co-transformed with *AoErg10B-DsRed* and *AoErg10C-DsRed* and PTS-GFP vectors. Left to right: fluorescent image of DsRed, GFP, merged image of DsRed and GFP. (B) The mycelium of *A. oryzae* 3.042 *△pyrG* transformed with AoErg10B^ΔMTS^-DsRed and AoErg10C^ΔMTS^-DsRed. Left to right: fluorescent image of DsRed, bright field, and merged image of DsRed and bright field.

Figure S4. The phenotype of *Pot1/erg10* transfromed *S. cerevisiae*. Growth of wildtype, *erg10* weak mutant (Y40985), *ScPot1*/WT and *ScPot1*/*erg10* transformants on YPD and YPG medium under 30°C and 37°C.

Figure S5. Phenotypes of *AoErg10s* over expression strains. (A) Scheme showing different transgenic strains. (B and C) Colony morphologies of the control (AoCK, wild-type *A. oryzae*), and AoErg10s over expression strains on the DPY and CD medium incubated at 30 °C for 72 h. Both DPY and CD medium use maltose as carbon source.

